# Spatial Meta-transcriptomes of human and murine intestines

**DOI:** 10.1101/2021.12.13.472336

**Authors:** Lin Lv, Ru Feng, Xue Li, Xiaofei Yu, GuoQiang Chen, Lei Chen

## Abstract

We developed an analysis pipeline that can extract microbial sequences from Spatial Transcriptomic data and assign taxonomic labels to them, generating a spatial microbial abundance matrix in addition to the default host expression one, enabling simultaneous analysis of host expression and microbial distribution. We applied it on both human and murine intestinal datasets and validated the spatial microbial abundance information with alternative assays. Finally, we present a few biological insights that can be gained from this novel data. In summary, this proof of concept work demonstrated the feasibility of Spatial Meta-transcriptomic analysis, and pave the way for future experimental optimization.

## Background

Spatial transcriptomic sequencing has revolutionized the biological research field. This class of technologies combine the strength of two pillars of modern biological research, sequencing and imaging. It generally works by capturing the messenger RNA from a permeabilized tissue slice, and label these RNA molecules with 2D spatial barcodes [1, 2]. Complex tissues such as the brain and tumor saw most utilization [3–9]. In the intestines and other organs, microbes live alongside or within close proximity to host cells and meta-genomic sequencing has long been employed to study the complex microbial composition on various host body sites. These studies, usually referred as microbiome study, have greatly improved our understanding of the human biology, microbes were fund in never-thought before places [10, 11] and showed intriguing dynamics [12]. However, these microbiome studies generally lacked spatial resolution and attempts at addressing this limitation is just emerging [13, 14]. Inspired by a recent study about COVID19 where virus sequences were recovered from host single cell transcriptomic data and analyzed alongside host data, we wanted to see if it is possible to capture microbial sequences in spatial transcriptomic sequencing and present our proof of concept work here.

## Results & Discussion

We performed spatial transcriptomic sequencing, using the Visium kit from 10X Genomics, on both human and murine intestinal samples, where microbial presence is well-known. The human samples included dissected colon samples from two colorectal cancer patients, samples from the tumor sites and histologically normal sites distant from the tumor were included. The murine samples came from both small intestine and large intestine, and cross-sectional slices from 3 mice were put in a single window (Fig. 1A). Each window was sequenced to at least 85k reads per covered spots, and the captured RNAs were on average 11k per spots, similar to that of published results [1].

**Fig. 1.**
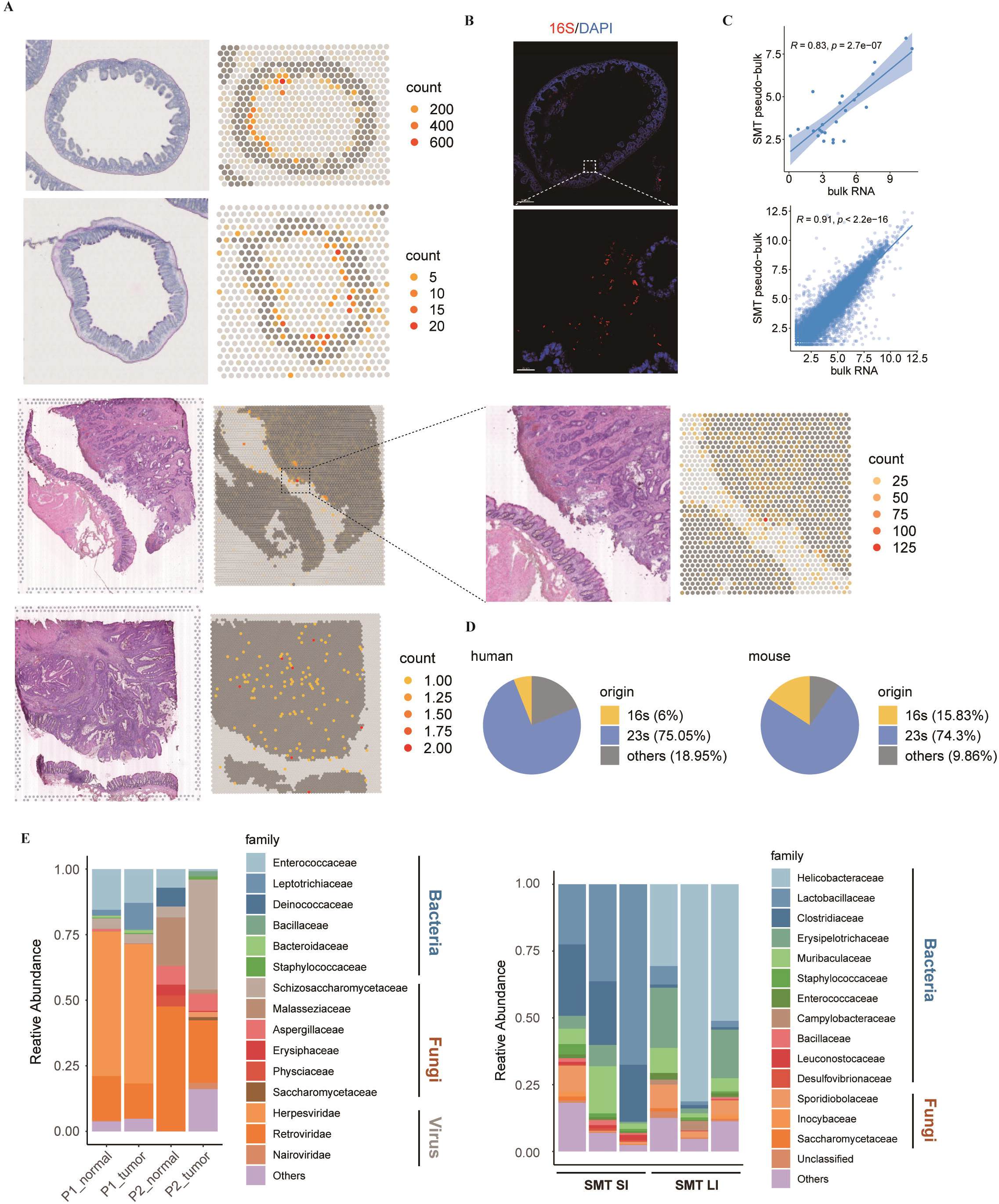
Spatial meta-transcriptomics demonstrating abundant microbial signal in mouse and human intestinal samples with appreciable diversity. **(A)** Feature plot showing abundance of microbial sequences in mouse small intestine (panel 1), mouse colon (panel2), human colonic normal and tumor samples (panel 3 and 4, with same design, in which normal and tumor samples occupy a single square area), colors indicate number of microbial sequences in form of Unique Molecular Identifier (UMI). **(B)** Small intestine slices from mice was stained with FISH probes against bacterial 16S rRNA. Boxed areas in the top row are magnified below. Scale bars, 300um (top), 20 μm (bottom). **(C)** Correlation analysis between SMT and bulk RNA sequencing data of mouse small intestine. Upper panel demonstrates correlation between microbial sequence count at family level, bottom panel shows correlation between host gene expression. **(D)** microbial read composition, data were normalized by samples. **(E)** Barplot illustrating composition of microbial sequences of human (upper) and mouse (bottom) samples at family level.

We developed an analysis pipeline to extract microbial information from the ST data. In short, unmapped reads were aligned against the NCBI NT database and the mapped reads in this round were subsequently assigned tax IDs. For reads coming from the same UMI (Unique Molecular Identifier) group, a common ancestor was called and assign to that UMI. The end result is a spatial species abundance matrix produced alongside the spatial host gene expression matrix generated via the standard spatial transcriptomics analysis.

The percentage of microbial sequences against total reads varied across different samples, ranging from 7.32 × 10^-7 in human samples to 2.26 × 10^-3 in murine ones. The Visium kit used poly T to capture messenger RNA and in theory microbial sequences without polyA tail would be depleted. While this is indeed the case, some spurious random priming did occur and sequences without polyA were still captured. This is in line with report from the RNA-velocity paper[15–17], where immature messenger RNA, which lacked polyA tail, account for about 25% of the data in 10X Genomics Chromium single cell platform, which uses polyA capture. Furthermore, ribosomal RNAs are the most abundant RNA category in a cell, account for more than 80% of total RNA. Ribosomal RNA is also the molecular marker of choice for microbial taxonomy. Accordingly, the majority of the microbial sequences recovered in SMT were from ribosomal RNAs (Fig. 1D). For these reasons, the microbial signals obtained from our Spatial Meta-Transcripotomic (SMT) analysis was unlikely to be merely artifacts.

For comparison, we applied our microbial sequence extraction method on single cell sequencing data derived from un-sorted intestinal samples and the microbial signal thus produced were very likely drenched in noise as it was relatively evenly spread across different cell types (Fig S1A). This may have to do with the fact that only intra-cellular bacteria can be captured in these conditions and these bacteria were rare and that the gentle digestive environment inside a droplet was ineffective for most microbes.

To further evaluate the extent of contaminating microbial sequences during experiments versus true organ resident microbial signals. We plotted the microbial abundance of all spots, regardless of whether it’s covered by tissue or not (Fig 1A). For both human and mouse, the luminal side of the intestines harbored more microbial signal as is expected. The human tumor tissue also contained higher abundance of microbes and also more microbes penetration deeper into the tissue, as would be expected from the compromised barrier function in tumor. These clear and expected patterns gave validation to our SMT methods as noise signal would show up relatively randomly across the capture slide.

To compare results with alternative methods, for the mice samples, we visualized slices close in proximity from the same embedded sample by Florescent In Situ Hybridization (FISH) with the bacterial probe EUB338 (Fig 1B). The FISH result was in agreement with SMT with bacteria presence concentrated in the intestinal lumen. Of the three slices shown in the same window, the general bacteria abundance varies and corresponds well with SMT result. In another word, the intestine piece with most bacteria as shown by FISH is the one with most bacteria sequences in SMT result. These variances could be due to variance in processing individual sample and in difference in exact sample origin in the gut, i.e., more distal or proximal. All these observations lend credibility to SMT approach.

After the assessment of contamination, we continued to evaluate the biases in the microbial signals collected in SMT. Because during the Visium process, the host tissue slides were permeabilized to release its RNA and this process is far from ideal for microbial sequence capture. This brought up the question if certain microbial species would be more amicable to this condition and the final result would present a biased picture of the true microbiome. We also cautioned that if the capture efficiency of microbial sequences were so low as to make the result subject to great fluctuation that will also make the result unreliable. To answer these questions, we extracted RNAs from corresponding tissue slices and performed total RNA sequencing, in which polyA based enrichment is not used and only host ribosomal RNA is depleted. Similar to our SMT pipeline, we processed the bulk total RNA-Seq data to generate microbial abundances, one abundance profile for on slice. To compare with bulk sequencing data, we then combined the SMT’s per spot abundance profiles for one window into pseudo-bulk ones and first evaluated the correlation between host genes and then microbial abundances at family level (Fig. 1C). The results showed that the host gene correlation was at 0.91 with p value 2.2 × 10^-16 and that the microbial abundance reached 0.83 with p value of 2.7 × 10^-7, comparable to host gene levels and thus validating SMT approach.

Due to its RNA-Seq nature, our SMT methodology captures Fungi and viral sequences as well as bacteria ones. Zoonotic viruses generally have mRNA that resemble their hosts and interestingly we can identify one such virus infection case in great spatial detail in one of our human CRC datasets. In one out of the two patients examined, viral sequences reaching as high as 3.4% were seen in some spots at the tumor site. This virus, identified as Cytomegalovirus, is known to infect fibroblast, and a deconvolution of the infected spots, which were all 55μm in diameter and thus contained many cells, showed them to be mostly fibroblast (Fig 2A). We next asked what the immediate consequence of viral infection was on these fibroblasts. To answer this, infected spots were compared to un-infected fibroblast rich spots. Interestingly, among the differentially expressed genes recovered, interferon related genes were not found, potentially due to the immune-compromised nature of the tumor micro-environment. Still, some known genes were found, including HLA-E which is know to be up-regulated by cytomegalovirus infection, IL32 and TNIP, both upregulated and involved in host defense (Fig 2B).

**Fig. 2.**
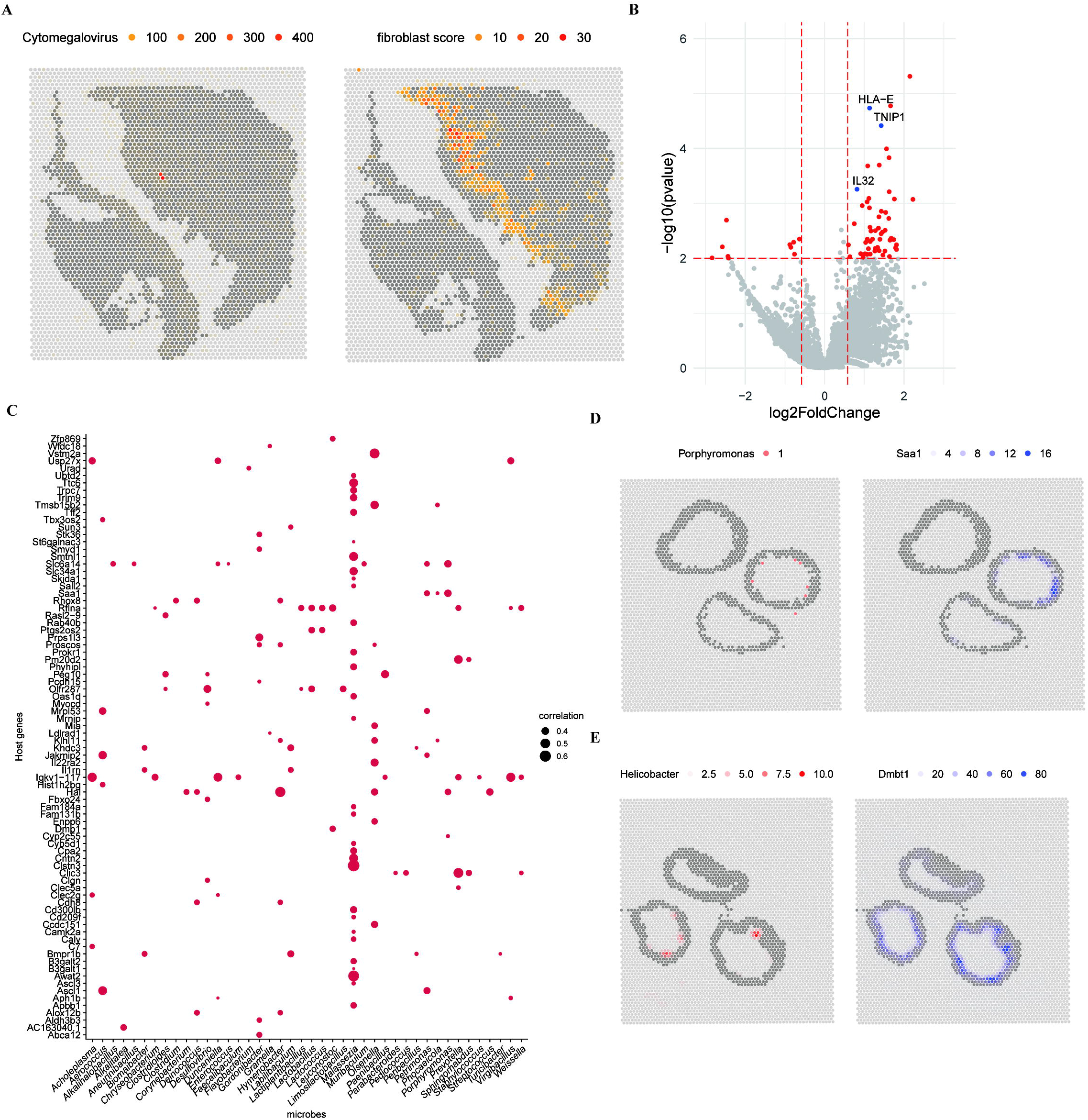
Spatial meta-transcriptomics reveal novel and commonly recognized host-microbe interactions. **(A)** Feature plot showing abundance of sequence from genus *Cytomegalovirus* in human colorectal cancer sample (left) and module score of fibroblasts (right). **(B)** Scatter plot of differential expression analysis with cytomegalovirus enriched spots and cytomegalovirus absent spots that were tagged with high fibroblast score. Genes with high significance value (<0.001) were presented with red color, genes responsible for antiviral response in those genes were labeled blue. **(C)** Dot plot showing colocalized correlation between host gene expression and genus level microbial abundance in mouse small intestine. **(D)** Spatial feature plot showing *Porphyromonas* enrichment and Saa1 expression. **(E)** Spatial feature plot showing *Helicobacter* enrichment and Dmbt1 expression.

The local cellular response to infection shown above is a good testament to SMT’s capability in investigating microbe-host interaction. Compared to other methods in this context, FISH based for example, SMT’s strength comes in its taxonomical resolution and systemic nature. For a demonstration, we used the murine dataset as it included multiple individual animals and covered complete cross-section of the intestines. We asked, which host genes and which microbes showed great spatial correlation. Due to the sparse nature of the microbial signals and to lesser degree, some of the host genes, we first smoothed both categories of signals (Fig. S2A, Methods). Then spatial correlation analysis was performed and revealed a number of interesting interactions (Fig 2C for small intestine, Fig S2B for large intestine). Among the most significant, the bacteria *Porphyromonas* showed high correlation with a series of immune defense related genes such as Saa1 in the small intestine, while *Helicobacter* correlated with Dmnt1 in the large intestine, and consistently, Helicobacter also correlated with Saa1(Fig 2B, Fig S2B). We can also highlight individual interactions by plotting the intensities of involved host gene and microbe on the slice and directly visualize the extent of the interaction, whole sections of intestines in this particular case (Fig 2DE, Fig S2C).

Our current SMT method generates spatial microbial abundance matrix alongside the spatial host expression matrix. The underlying spatial transcriptomic methods are rapidly evolving[18], reaching higher resolution and obtaining more sequences per area, translating to higher sensitivity. SMT too will correspondingly reap the benefit. Furthermore, by simply capturing more sequences or by using specifically designed capture oligos, a bacteria specific 16S capture oligo alongside the polyT capture oligo for example, it may become possible to actually profile the microbial transcriptomes. For tissues where microbes were abundant such as the intestines, SMT enables the systemic study of host and microbe interaction. In other more sterile tissues, SMT will shed light on the consequence of microbial presence, for example, how microbes travel to remote tumor site (as in non-intestinal solid cancers) and help settle the debate on the whether certain body site is sterile or not, i.e., in-utero fetus[19, 20].

## Conclusions

Our proof of concept work demonstrated the feasibility of Spatial Meta-Transcriptomic sequencing and analysis. Our analysis framework already extracted spatial microbial abundance information out of data generated by currently commercially available kits. We further demonstrated that true signals outweigh contaminations and that the biases are low. Actually, the FISH result suggested the current SMT method to be very sensitive. Our work paves the way for future development of this technology, which will further increase sensitivity of SMT and its taxonomical resolution. Such methods will enable simultaneous analysis of host expression with resident microbial abundances, and study host and microbial interactions in never-before seen resolution and doing so in a systematic manner.

## Methods

### Human/Murine sample collection and processing

We collected samples of pathologically diagnosed with CRC from Rui-jin hospital, Shanghai Jiao Tong University. Tissue samples were embedded in optimal cutting temperature compound and stored at −80°C. Before the tissue optimization experiment was performed, the RNA quality was checked (RIN>7.0). The tumors are resectable and discussion by clinicians. Six-week-old male C57BL/6 mice were ordered from Shanghai SLAC Laboratory Animal Co., Ltd. Then maintained in SPF experimental animal center of Fudan University. For antibiotics treatment, six-week-old mice were orally gavaged with 0.5mg/mL vancomycin (RPI),1mg/mL metronidazole, 1mg/mL ampicillin, 1mg/mL neomycin, 1mg/mL gentamycin dissolved in 1XDPBS for one week. The maximum number of adult mice in a cage is 5. Feces and serum were collected and stored at −80°C. Tissue samples(intestine and colon) were embedded in optimal cutting temperature compound and stored at −80°C, the RNA quality was checked(RIN > 7.0).

### Spatial transcriptomics

Tissues were cut into 10μm sections and processed using the Visium Spatial Gene Expression Kit (10x Genomics) according to the kit’s instructions. First step, CRC tissue, mice tissue permeabilization condition was optimized using the Visium Spatial Tissue Optimization Kit, which was 18 min in mice and 28 min in human found to be maximum fluorescence signal in both tumor and normal regions. In detail, 4 samples from 2 patients were sequenced by ST. Then were stained with H&E and imaged using a Leica DM6000 microscope under a 20X lens magnification. Next, reverse transcription, second strand synthesis & denaturation, cDNA amplification & QC, Visium spatial gene expression library construction was following the manufacturer’s instructions. The resulting complementary DNA library was checked for quality control, then sequenced using an Illumina NovaSeq 6000 system.

### Spatial transcriptomic sequencing and data processing

Raw reads from 10x Genomics Visium spatial sequencing were aligned to the human transcriptome GRCh38-3.0.0 reference or mouse transcriptome mm10-3.0.0 reference using 10x Genomics SpaceRanger v.1.0.0 (https://support.10xgenomics.com/spatial-gene-expression/software/downloads/1.0/#spacerangertab) and exonic reads were used to produce mRNA count matrices for these samples. HE histology images were also aligned with mRNA capture spots using SpaceRanger.

### Spatial meta-transcriptomic analysis

An in-house pipeline was used for pre-processing of bam file generated by SpaceRanger. Unmapped reads were extracted from bam file and filtered for non-polynucleotide sequence in preparation for subsequent alignment. After that, unmapped reads were aligned against nt database with blastn 2.10.1+, the blast output were formatted to preserve query ID, taxID, subject title, alignment length, e-value, identity, coverage as well as UMI and spatial barcode which were extracted from bam file. Taxa levels from kingdom to species were called with taxID by querying a taxa table from NIH taxonomy online port. UMI counts of certain taxID in a single spot were called by counting unique UMI sequences belonging to respective spatial barcode and thus we could generate a count matrix with taxID and spatial barcode as row names and column names respectively.

The UMI count matrix were then imported into R to generate a Seurat object (Seurat 4.0.1)[21] with HE images that had been included in SpaceRanger output directory. Simultaneously, the count matrix and taxa level info were also used to create a phyloseq object (phyloseq 1.32.0)[22] with an applied taxa level and integrated into the assay slot of Seurat object.

### FISH

OCT tissue slides were fixed with methyl alcohol at −20 °C. Slides were stained using the FISH kit (Guangzhou Exon) following manufacturer s protocol. Cy3 labelled Probes (EUB338 - GCTGCCTCCCGTAGGAGT) were hybridized overnight at 37°C. Sections were washed at room temperature and 60 °C in washing buffer1 for 5 minutes successively, followed by two washed at 37 °C for 5 minutes in washing buffer2. Staining was visualized with the Leica TCS SP8 confocal microscope at 20X and 63X. The images were edited using Imaris Cell Imaging Software.

### Bulk total RNA-Seq

OCT tissue slides were used for total RNA isolation with TRIzol (Invitrogen) and subjected to RNA sequencing using Illumina NextSeq 500 system (75-bp paired-end reads). The raw reads were aligned to the mouse reference genome (version mm10) and human reference genome (version hg38) using HISAT2 RNA-sequencing alignment software[23]. The alignment files were processed to generate read counts for genes using SAMtools[24] and HTSeq[25]. Read counts were normalized and transmitted to differential analysis using R package DESeq2[26]. P values obtained from multiple tests were adjusted using the Benjamini-Hochberg correction. Gene ontology consortium (GO) and Kyoto Encyclopedia of Genes and Genomes (KEGG) pathway analysis was performed using R package clusterProfiler[27].

## Data & Code Availability

The raw data of mouse study, analysis pipeline and the matrix for both human and mouse will be made available upon publication.

## Acknowledgements

We would like to thank the Rui-jin hospital of Shanghai for their help with tumor tissue preparation and specimen acquisition. We would also like to thank the sequencing diagnosis center of Shanghai Institute of immunology for their services. This work was in part supported by the School of Life Sciences Fudan university.

## Figure legend

**Fig. S1 Microbiome composition and abundance cross samples. (A)** UMAP visualization of the un-sorted human colorectal sample, showing the formation of 11 main clusters. **(B)** Microbial signals with high abundance (>0.02%) illustrated in U-MAP plot. **(C)** Microbiome composition cross SMT and bulk RNA seq data at kingdom level. **(D)** Microbiome composition in bulk RNA seq data at family level.

**Fig. S2 Host-microbe interaction analysis with SMT data. (A)** Feature plot of original UMI count and smoothed UMI count with same ST data showing appreciable effect of smoothing. **(B)** Dot plot showing co-localized correlation between host gene expression and genus level microbial abundance in mouse colon. **(C)** Spatial feature plot showing Helicobacter enrichment and Saa1 expression.

