## Supplementary figures and images for "Spatial Meta-transcriptomes of human and murine intestines"

### Fig. S1

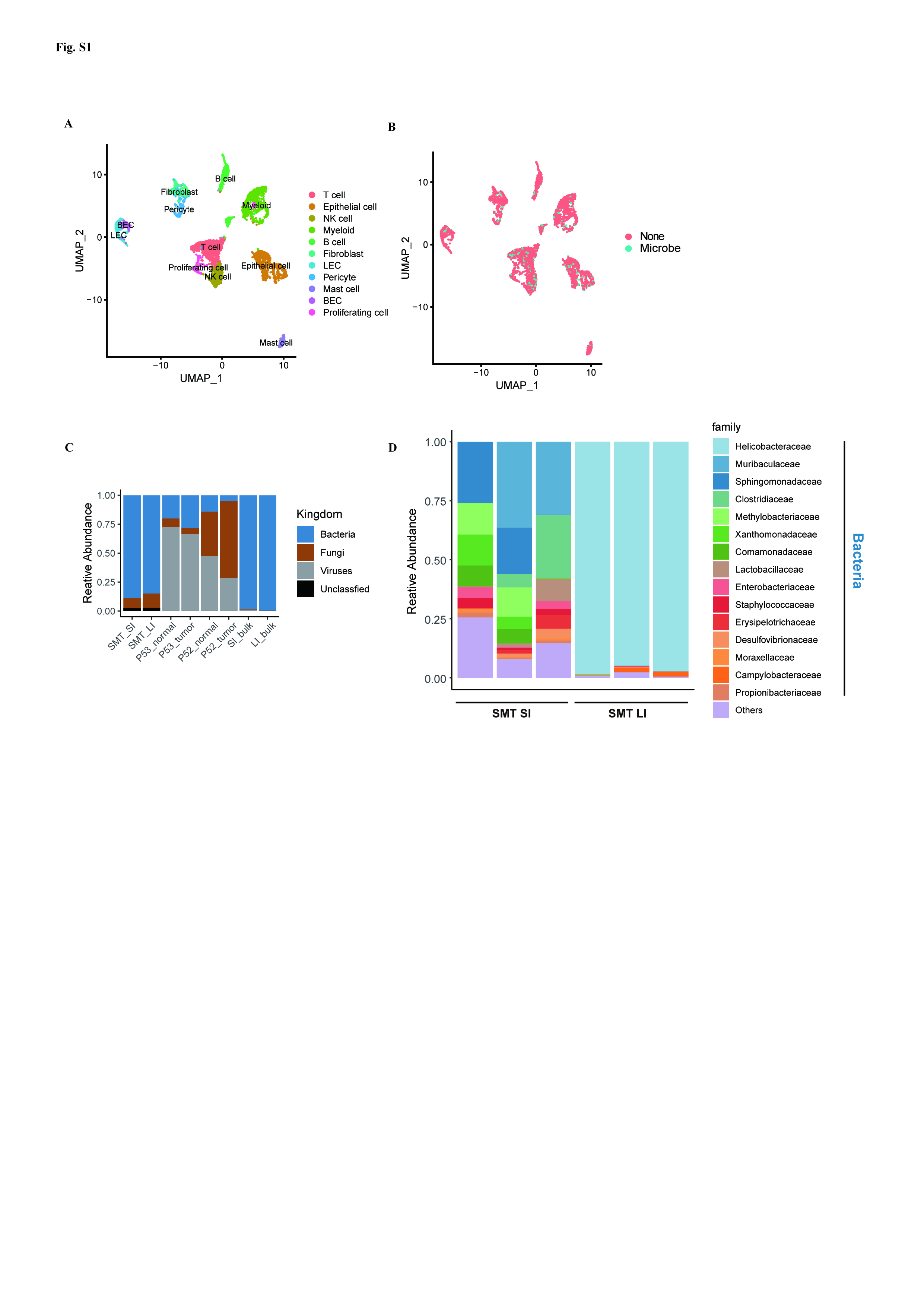

### Fig. S2

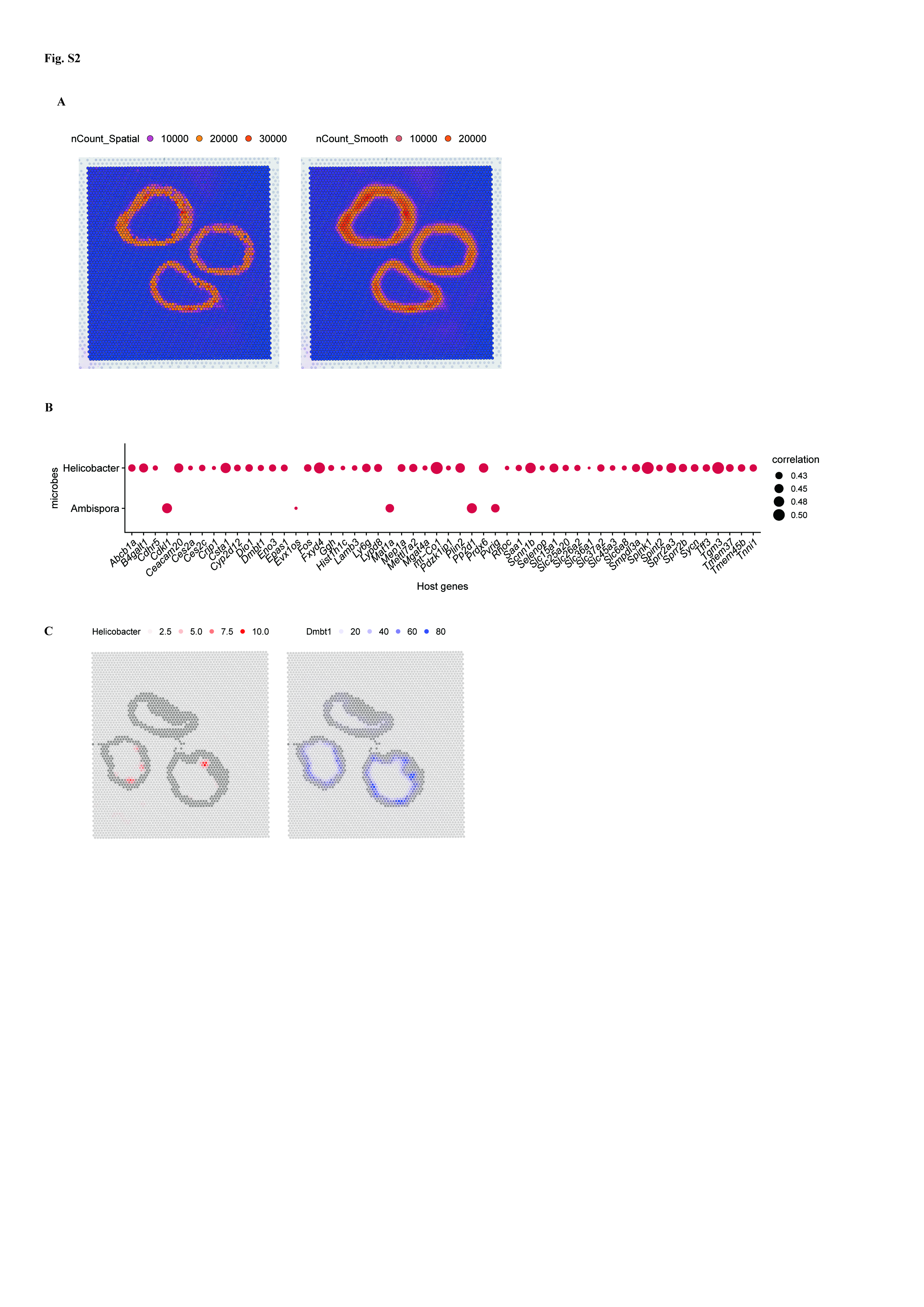
